# Two distinct actin filament populations have effects on mitochondria, with differences in stimuli and assembly factors

**DOI:** 10.1101/701771

**Authors:** Tak Shun Fung, Wei-Ke Ji, Henry N. Higgs, Rajarshi Chakrabarti

## Abstract

Recent studies show that mitochondria and actin filaments work together in two contexts: 1) increased cytoplasmic calcium induces cytoplasmic actin polymerization that stimulates mitochondrial fission, and 2) mitochondrial depolarization causes actin assembly around mitochondria, with roles in mitophagy. It is unclear whether these two processes utilize similar actin assembly mechanisms. Here, we show that these are distinct actin assembly mechanisms in the acute phase after treatment (<10 min). Calcium-induced actin assembly is INF2-dependent and Arp2/3 complex-independent, whereas depolarization-induced actin assembly is Arp2/3 complex-dependent and INF2-independent. The two types of actin polymerization are morphologically distinct, with calcium-induced filaments throughout the cytosol and depolarization-induced filaments as “clouds” around depolarized mitochondria. We have previously shown that calcium-induced actin stimulates increases in both mitochondrial calcium and recruitment of the dynamin GTPase Drp1. In contrast, depolarization-induced actin is temporally-associated with extensive mitochondrial dynamics that do not result in mitochondrial fission, but in circularization of the inner mitochondrial membrane (IMM). These dynamics are dependent upon the protease Oma1 and independent of Drp1. Actin cloud inhibition causes increased IMM circularization, suggesting that actin clouds limit these dynamics.

**Summary statement:** Mitochondrial depolarization induces Arp2/3 complex-dependent actin clouds that restrain mitochondrial shape changes induced by Oma1 on the inner mitochondrial membrane. A distinct actin network stimulates mitochondrial fission in response to calcium.

## Introduction

Mitochondria have traditionally been viewed as energy-generating organelles, through oxidation of metabolic substrates and creation of a proton gradient across the inner mitochondrial membrane (IMM), with the subsequent passage of protons back into the matrix being coupled to ATP synthesis (Kennedy and Lehninger, 1949; Pagliarini and Rutter, 2013). However, it is increasingly clear that mitochondria communicate frequently with the rest of the cell and are therefore important signaling organelles. For example, mitochondrial release of cytochrome c triggers cell death (Liu et al., 1996), mitochondrially-generated reactive oxygen species activate hypoxia-related genes (Al-Mehdi et al., 2012; Chandel et al., 1998) and mitochondrial heat shock proteins promote cytosolic calcium-mediated signaling (Biswas et al., 1999; Martinus et al., 1996). Mitochondria also participate in innate immunity by serving as platforms for downstream signaling to facilitate anti-microbial host cell responses (West et al., 2011). Finally, changes in the IMM proton gradient have immediate signaling effects, with IMM depolarization causing stabilization of the PINK1 protein kinase, whose downstream targets include the PARKIN E3 ubiquitin ligase (Pickles et al., 2018). Mitochondrial depolarization also activates an IMM protease, OMA-1, which proteolytically cleaves the dynamin-family GTPase Opa1 (Ehses et al., 2009; Head et al., 2009; Ishihara et al., 2006).

A growing number of studies suggests that actin polymerization participates in mitochondrial communication and dynamics. During apoptosis, both a C-terminal actin fragment (Utsumi et al., 2003) and the actin-binding protein cofilin translocate to mitochondria, with evidence that translocation of active cofilin is important for downstream cytochrome c release and apoptotic response (Chua et al., 2003). Inhibition of ATP synthase by oligomycin results in mitochondrial fission, which is attenuated by the actin polymerization inhibitors cytochalasin D or latrunculin A (De Vos et al., 2005). In another study, actin and myosin II were shown to play a role in translocation of the dynamin GTPase Drp1 to mitochondria, resulting in mitochondrial fission (DuBoff et al., 2012).

We have previously found that elevated cytosolic calcium activates the endoplasmic reticulum (ER)-bound formin INF2, which stimulates actin polymerization that leads to mitochondrial fission (Chakrabarti et al., 2018; Hatch et al., 2016; Ji et al., 2017; Ji et al., 2015; Korobova et al., 2013). INF2-mediated actin polymerization stimulates constriction of both mitochondrial membranes during fission: IMM constriction is enhanced by increased calcium transfer from ER to mitochondrion, and OMM constriction is enhanced by increased Drp1 recruitment. This pathway also requires non-muscle myosin II (Chakrabarti et al., 2018; Korobova et al., 2014), and the mitochondrially-bound SPIRE 1C protein might also participate (Manor et al., 2015).

A somewhat different type of mitochondrially-associated actin polymerization has been reported in several studies. Dissipation of the mitochondrial proton gradient using the uncoupler FCCP causes rapid accumulation of an extensive cloud of actin filaments around depolarized mitochondria (Li et al., 2015). Similar actin clouds were observed in both unstimulated cells and cells treated with the uncoupler CCCP and were shown to be dependent on both Arp2/3 complex and formin activity (Kruppa et al., 2018; Moore et al., 2016). On a similar time scale as actin cloud formation, mitochondria become less elongated, consistent with an increase in mitochondrial fission (Li et al., 2015; Moore et al., 2016). At a later stage after mitochondrial depolarization, a second wave of actin polymerization encircles depolarized mitochondria, and is proposed to prevent their fusion with other mitochondria in a myosin VI-dependent manner (Kruppa et al., 2018).

These studies raise a question concerning actin polymerization and mitochondrial function. Are there multiple ways in which actin interacts with mitochondria, or do these studies represent variations on a common actin polymerization pathway? Our present work attempts to clarify this issue. We show that calcium-induced actin polymerization is INF2-dependent and Arp2/3-independent, whereas depolarization-induced actin polymerization is Arp2/3-dependent and INF2-independent. Calcium-induced actin polymerization is significantly faster than depolarization-induced actin polymerization and is less tightly associated with mitochondria. Spontaneous mitochondrial depolarization causes Arp2/3-dependent actin polymerization around the depolarized mitochondria similar to that of CCCP-induced actin filaments. While mitochondrial depolarization results in extensive mitochondrial shape changes on a similar time course to actin polymerization, these shape changes are not dependent on actin polymerization or Arp2/3 complex. In fact, the shape changes are due to IMM dynamics and are dependent on the IMM protease OMA1. Inhibition of actin cloud assembly causes an increase in CCCP-induced mitochondrial shape changes, suggesting that actin clouds inhibit these shape changes. In summary, we show that two distinct types of actin filaments, differing in morphology and assembly mechanisms, have differing effects on mitochondria.

## Results

### Distinct actin polymerization mechanisms induced by mitochondrial depolarization or cytoplasmic calcium

We compared the actin bursts stimulated by the calcium ionophore ionomycin with those induced by the mitochondrial depolarizer CCCP in U2OS cells, using live-cell imaging of actin filaments (GFP-F-tractin) and mitochondrial matrix (mito-BFP). In serum-containing medium, ionomycin treatment results in a transient increase in cytoplasmic calcium, with a T_1/2_ of <5 sec and a return to baseline within 2 min (Chakrabarti et al., 2018), while CCCP causes mitochondrial depolarization within 30 sec and persisting for at least 10 min (Figure S1A). Both stimuli induce transient actin polymerization responses that differ in morphology and kinetics. Morphologically, ionomycin-induced actin polymerization occurs throughout the cytosol, whereas CCCP-induced actin polymerization occurs as “clouds” that are closely associated with mitochondria (Figure 1A, Figure S1B). These features are best appreciated in a medial Z-plane, (Figure S1B, Movies 3, 4), due to less interference from basal actin stress fibers. However, actin polymerization in response to both stimuli is apparent at the basal surface as well (Figure 1A, Movies 1, 2). Kinetically, ionomycin-induced actin dynamics are more rapid than those induced by CCCP (Figure 1B), both in the polymerization (T_1/2_ actin polymerization values: 12.2 ± 5.9 sec and 133 ± 78.1 sec respectively) and depolymerization phases (T_1/2_: 29.4 ± 12.9 sec and 79.31 ± 38.9 sec respectively) (Figure S2A).

**Figure 1:**
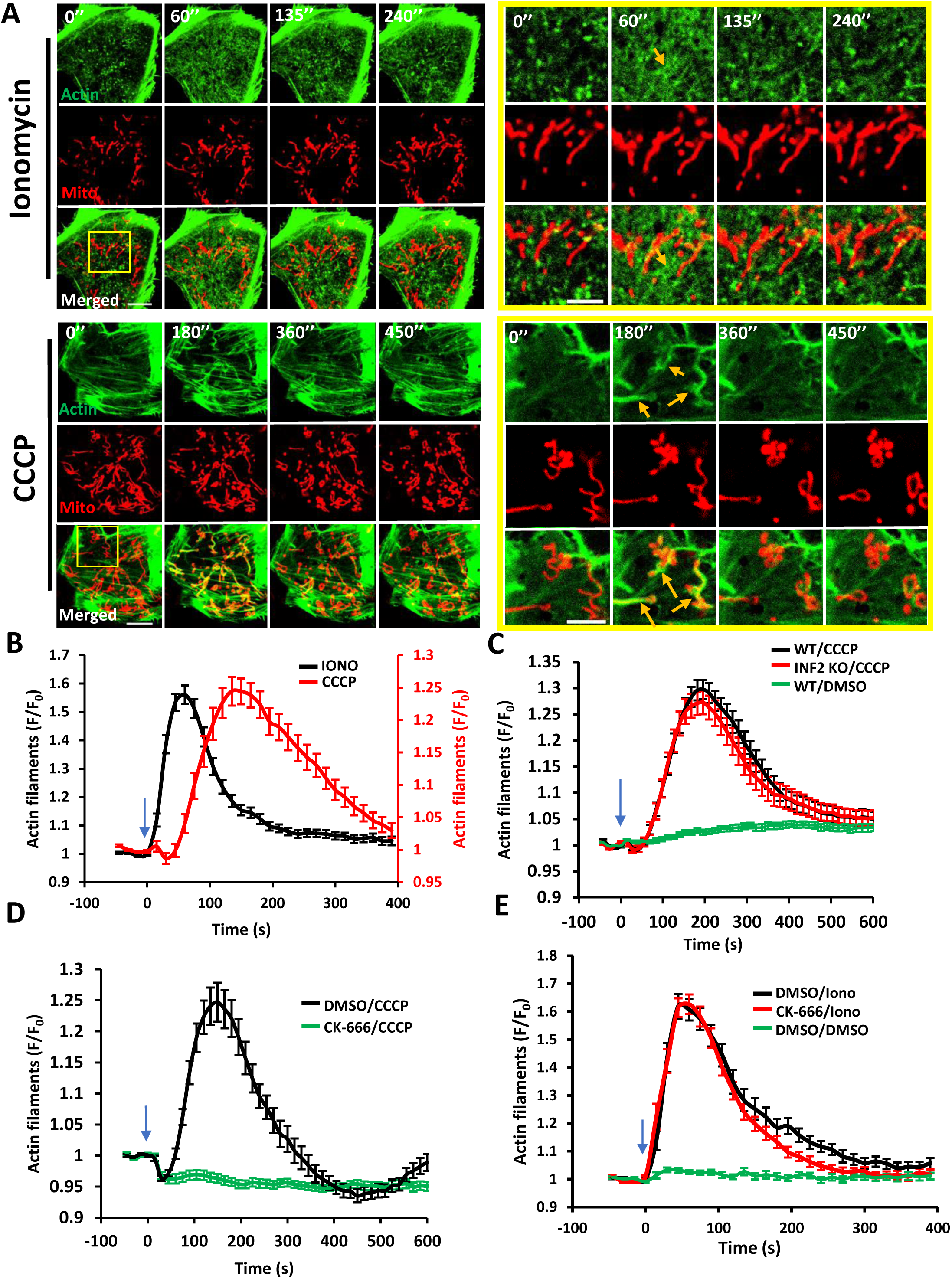
Distinct actin structures assemble in response to two stimuli: increased cytoplasmic calcium and mitochondrial depolarization. **(A)** Time-lapse image montage of ionomycin-induced (top) and CCCP-induced (bottom) actin polymerization for U2OS cells transfected with GFP-F-tractin (green) and mito-BFP (red). Imaging conducted at the basal cell surface. Ionomycin or CCCP added at time point 0. Scale bar: 10μm. Inset scale bar: 5μm. Corresponds to Movies 1 and 2. Movies 3 and 4 show similar time course in medial cell section. Orange arrow shows actin assembly in both cases. **(B)** Comparison of ionomycin-induced and CCCP-induced actin polymerization time course for U2OS cells. Data from three experiments. N = 30 cells/60 ROIs for ionomycin (4 μM) treatment, 27 cells/27 ROIs for CCCP (20 μM) treatment. 15 sec intervals. Blue arrow denotes drug addition. Error bar, ±SEM. **(C)** CCCP-induced actin polymerization in U2OS-WT and U2OS-INF2-KO cells. Data from three experiments. N=35 cells for WT, 39 cells for INF2 KO and 35 cells for WT cells stimulated with DMSO. 14 sec intervals. Blue arrow denotes CCCP addition. Error bar, ±SEM. **(D)** Effect of Arp2/3 complex inhibition on CCCP-induced actin polymerization. U2OS cells were treated with either DMSO or 100μM CK-666 for 30 min, then stimulated with 20μM CCCP (blue arrow). Data from three experiments. N=35 cells/35 ROIs for DMSO/CCCP, 41/41 for CK-666/CCCP. 15 sec intervals. Error bar, ±SEM. **(E)** Effect of Arp2/3 complex inhibition on ionomycin-induced actin polymerization. U2OS cells were treated with either DMSO or 100μM CK-666 for 30 minutes, then stimulated with DMSO or 4μM ionomycin (blue arrow). Data from three experiments. N=23 cells/46 ROI for DMSO/Iono, 25/50 for CK-666/Iono and 20/40 for DMSO/DMSO. 15 sec intervals. Error bar, ±SEM.

We next examined the actin assembly factors required for ionomycin- and CCCP-induced actin polymerization. Past results have shown that ionomycin-induced actin polymerization requires the formin INF2 (Chakrabarti et al., 2018; Ji et al., 2015; Shao et al., 2015; Wales et al., 2016). However, INF2 is not required for CCCP-induced actin polymerization, tested using either CRISPR-mediated INF2 KO (Figure 1C) or siRNA-induced INF2 knock-down (Figure S2B, C). Control experiments show that ionomycin-induced actin polymerization is abolished in INF2 KO cells (Figure S2D). These results show that INF2 is not required for depolarization-induced actin polymerization.

Arp2/3 complex has been shown to contribute to mitochondrially-associated actin polymerization in HeLa cells in the absence of stimulation (Moore et al., 2016), and to actin polymerization that occurs after prolonged CCCP treatment (Kruppa et al., 2018). We used the Arp2/3 complex inhibitor CK666 to test Arp2/3 complex involvement in the rapid actin bursts induced by ionomycin and CCCP. A 30-min pre-treatment with CK666 abolishes the CCCP-induced actin burst (Figure 1D), while having no clear effect on the ionomycin-induced actin burst (Figure 1E).

These results show that actin polymerization induced by increased cytoplasmic calcium and by mitochondrial depolarization differ in three ways: kinetically, morphologically, and in nucleation mechanism. Calcium-induced actin polymerization is INF2-dependent and Arp2/3 complex-independent, whereas depolarization-induced actin polymerization is Arp2/3 complex-dependent and INF2-independent.

### Spontaneous mitochondrial depolarization triggers Arp2/3 complex-mediated actin clouds

Mitochondria periodically undergo transient depolarization in the absence of uncoupler treatment (Lee and Yoon, 2014). We asked whether actin polymerization faithfully accompanies such transient depolarization in U2OS cells, using the mitochondrial membrane potential marker tetramethyl rhodamine methyl ester (TMRE). Occasional loss of TMRE fluorescence occurs in sub-populations of mitochondria (Figure 2A, movie 5). These depolarization events are generally transient, having a mean duration of 143.7 ± 106.5 sec (Figure 2E). Actin polymerization accompanies the majority (87.3% ± 6.3) of depolarization events (Figure 2A-C, movie 5), with an appreciable lag between depolarization and actin polymerization, taking an average time of 129.8 ± 95.0 sec (Figure S2E). The actual polymerization T_1/2_ (measured from the first detectable polymerization) is 57.2 ± 54.1 sec (Figure S2F).

**Figure 2:**
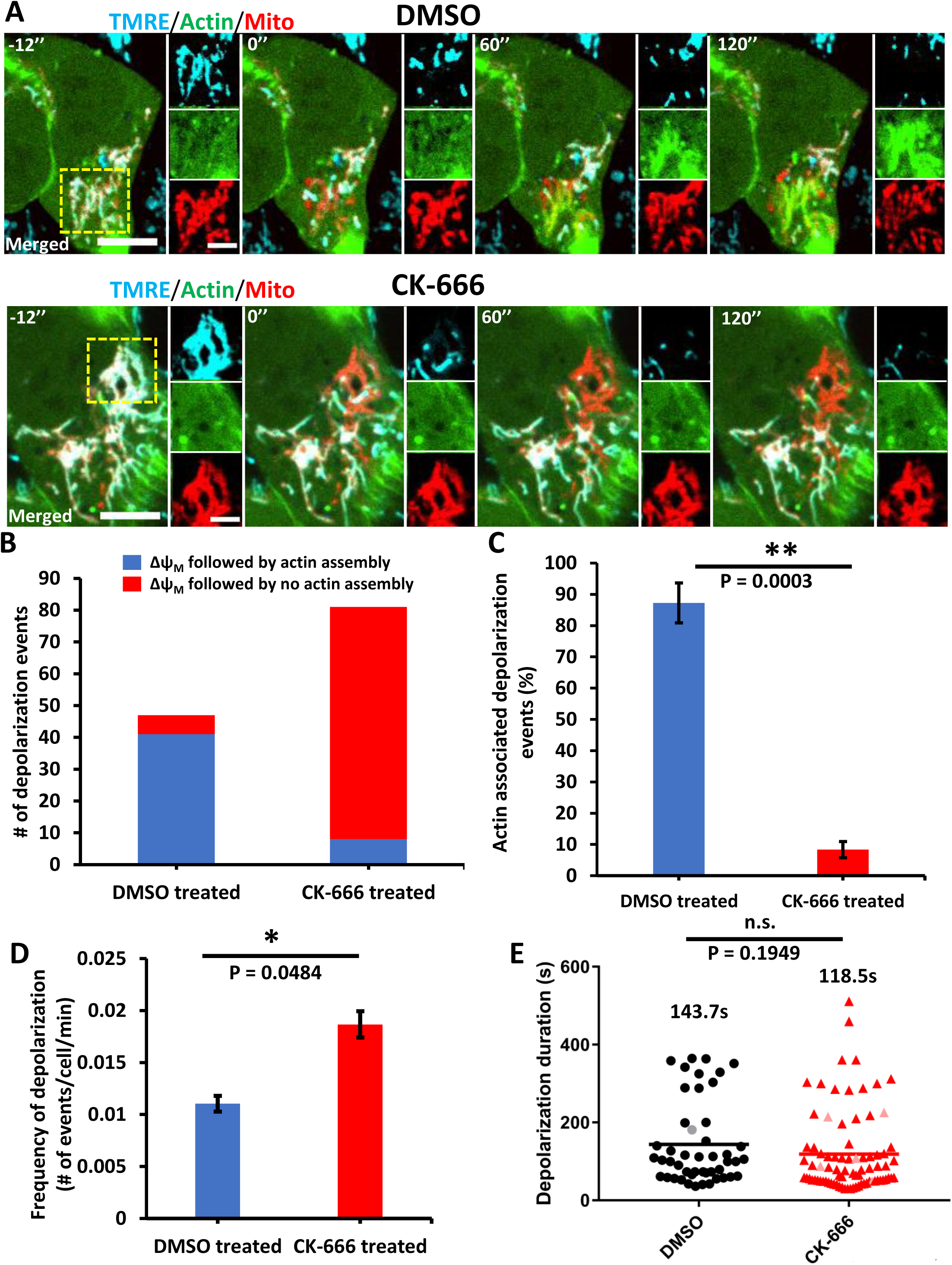
Actin polymerization accompanying transient depolarization of a sub-set of mitochondria in the absence of CCCP. **(A)**Time-lapse image montages of changes in mitochondrial polarization (TMRE, blue) and actin polymerization (GFP-F-tractin, green) in U2OS cells in the absence of uncoupler treatment. A mitochondrial matrix marker (Mito-BFP, red) is also included. Top panels show a control cell (DMSO) and bottom panels show a CK-666-treated cell (100 μM CK-666 for 30 min before imaging, and 50μM during imaging). Time 0 denotes start of a depolarization event. Panels to right of each time point denote zooms of boxed regions. Scale bars:10μm (5 μm for inset). Corresponds to Movies 5, 6. **(B)** Graph of spontaneous depolarization events associated with actin polymerization, in either control (DMSO) or CK-666-treated cells. Total numbers from 3 experiments (20 min imaging per cell, 1.2 sec intervals): DMSO, 213 cells, 47 depolarization events (41 events accompanied by actin assembly); CK-666, 217 cells, 81 depolarization events (8 events accompanied by actin assembly). **(C)** Graph of % depolarization events accompanied by actin polymerization for DMSO versus CK-666 treatment, from same data as panel B. Student’s unpaired t-test: p = 0.0003. Error bar: ±SEM. **(D)** Graph of depolarization frequency for DMSO versus CK-666 treatment, from data set described in panel B. Student’s unpaired t-test: p = 0.0484. Error bar: ±SEM. **(E)** Scatter plot of depolarization duration, from data set described in panel B. Mean depolarization durations: DMSO group (black line), 143.7 sec ± 106.5 sec; CK-666 group (red line), 118.5 sec ± 104.4 sec (standard deviation). Gray and pink points represent depolarization events that occurred after 16 minutes of imaging time and failed to repolarize at the end of the 20minutes imaging period. Student’s unpaired t-test: p = 0.1949.

We asked whether Arp2/3 complex is involved in actin polymerization induced by spontaneous mitochondrial depolarization. To this end, we pre-treated U2OS cells with CK666 for 30 min prior to imaging. Spontaneous depolarization of sub-populations of mitochondria still occurs in CK666 pretreated cells, similar to control cells (Figure 2A, movie 6). In fact, the frequency of spontaneous depolarization events is somewhat higher in CK666-treated cells (Figure 2D), while the average duration of depolarization events is not significantly different (Figure 2E). CK666 treatment, however, strongly reduces the actin polymerization events that occur after spontaneous depolarization (Figure 2A-C). These results further support the finding that the actin polymerization occurring around depolarized mitochondria is mediated by Arp2/3 complex in U2OS cells.

### Actin polymerization is not required for depolarization-induced mitochondrial shape change

In addition to inducing actin polymerization, CCCP-triggered mitochondrial depolarization induces rapid mitochondrial shape changes (Figure 1A). While these shape changes have often been described as mitochondrial fragmentation (Fu and Lippincott-Schwartz, 2018; Kwon et al., 2017), there is also evidence for other changes such as circularization (De Vos et al., 2005; Liu and Hajnoczky, 2011; Miyazono et al., 2018). Using the mitochondrial matrix marker mito-BFP in live-cell microscopy, we observe a variety of mitochondrial shape changes within the first 20 min after CCCP-induced mitochondrial depolarization. The major types of mitochondrial reengagements that we observe are 1) longitudinal splitting and 2) curling. While curling events typically occur at one end of the mitochondrion, splitting events can take place anywhere along the mitochondrion: at the ends, at the center, or at branch junctions (Fig 3A, movie 7). The end result of splitting or curling is circularization of the mitochondrion.

**Figure 3:**
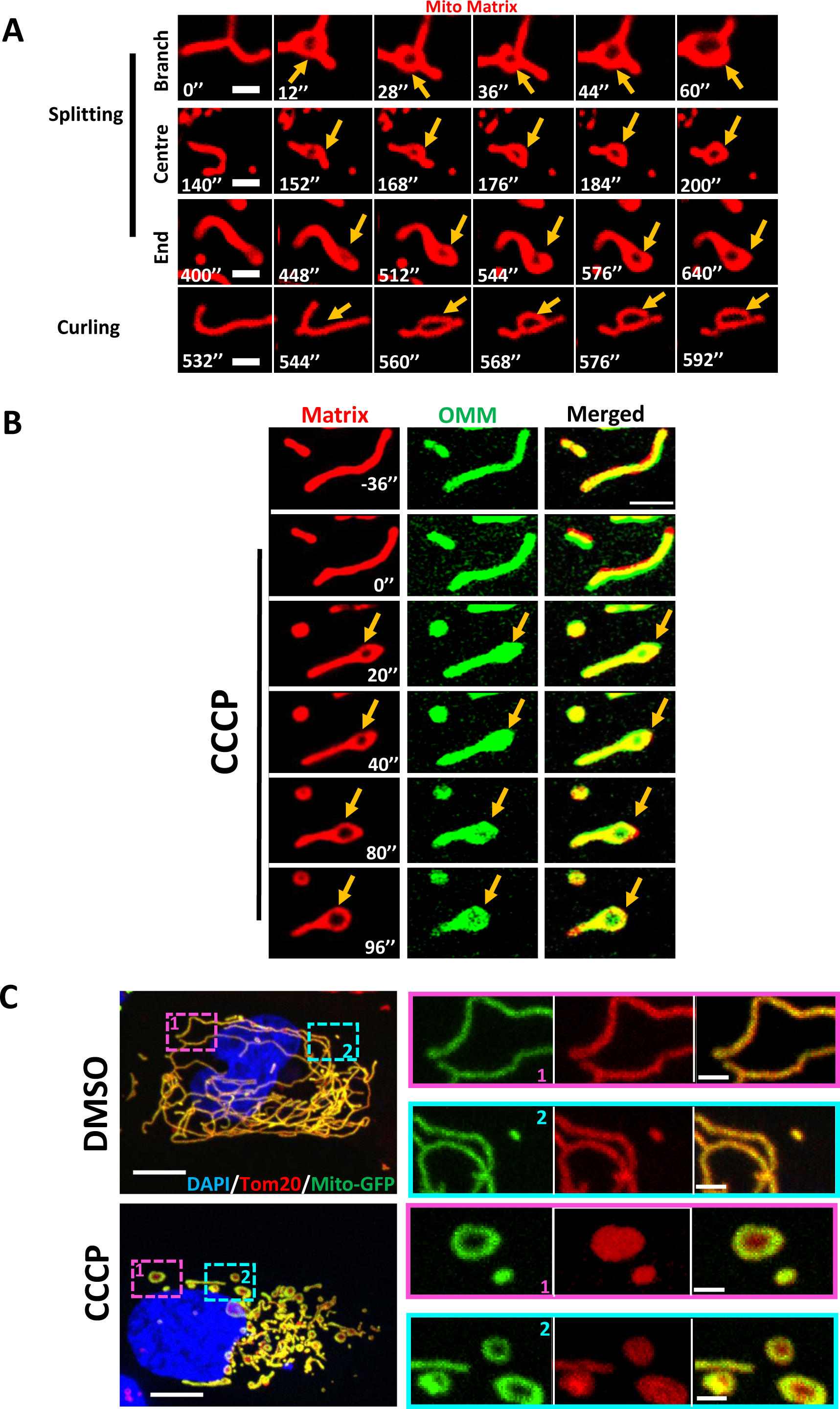
Depolarization-induced mitochondrial shape changes in U2OS cells. **(A)** Examples of mitochondrial matrix dynamics after depolarization. U2OS cells were transfected with the mitochondrial matrix marker mito-DsRed and treated with 20μM CCCP at time point 0. Confocal imaging was conducted at the basal region of the cell at 4 sec intervals, starting 10 frames before CCCP treatment. Scale bar: 2μm. Corresponds to Movie 7. **(B)** Dynamics of the OMM and mitochondrial matrix upon depolarization. U2OS cell transfected with Mito-DsRed (red) and Tom20-GFP (green) was treated with 20μM CCCP at time point 0. Airyscan images (basal region) were acquired at 4 sec intervals starting 10 frames before CCCP treatment. Scale bar: 2.5μm. Corresponds to Movie 8. **(C)** Maximum intensity projections of glutaraldehyde-fixed U2OS cells (transfected with GFP-Mito, green) after treatment with either DMSO (top) or 20μM CCCP (bottom) for 20 min. Cells were stained with anti-Tom20 (OMM, red) and DAPI (nucleus, blue). Z stacks were taken at step size of 0.4μm. Zooms show representative examples of mitochondrial circularization after CCCP treatment. Scale bars: 10μm and 2μm (inset).

We also used dual-color live-cell imaging to observe the dynamics of both the OMM (Tom20-GFP) and the matrix (Mito-dsRed) in the same cell. Interestingly, the matrix marker undergoes circularization, while the OMM marker remains intact across the center of this circularized region (Figure 3B, movie 8). This result is consistent in all cases analyzed (Figure S3A) and is similar to those previously reported by others (Miyazono et al., 2018), suggest that the IMM is the primary membrane undergoing rearrangement during depolarization-induced mitochondrial circularization.

To verify this result, we used fixed cell microscopy, first fixing cells with glutaraldehyde. Upon CCCP treatment for 20 min, numerous circular mitochondria are observed (Figure 3C). The mitochondrial matrix marker (transfected mito-GFP) displays a characteristic hollow donut shape, while the OMM marker (anti-Tom20 is intact throughout the circle. A similar pattern occurs upon formaldehyde fixation, although OMM staining is less regular in this case (Figure S3B). Using an antibody against the beta subunit of ATP synthase shows a similar result, although the ATP synthase staining is more irregular than the GFP matrix protein (Figure S3C).

We asked whether actin polymerization is required for these mitochondrial shape changes, quantifying shape change as the number of circular matrix structures (‘centroids’) present per overall mitochondrial area at specific timepoints after CCCP addition. Interestingly, while the actin sequestering drug latrunculin A (LatA) eliminates CCCP-induced actin polymerization (Figure 4A), it does not inhibit mitochondrial rearrangement (Figure 4B, 4C). In fact, the number of CCCP-induced centroids is increased by LatA treatment. Similarly, Arp2/3 complex inhibition by CK666 also increases CCCP-induced mitochondrial shape change (Figure 4D, 4E, Movie 9). CCCP-induced mitochondrial shape change also occurs in INF2-KO cells (Figure S3D, Movie 10). These results suggest that actin polymerization is not required for the acute mitochondrial shape changes that occur upon CCCP treatment.

**Figure 4:**
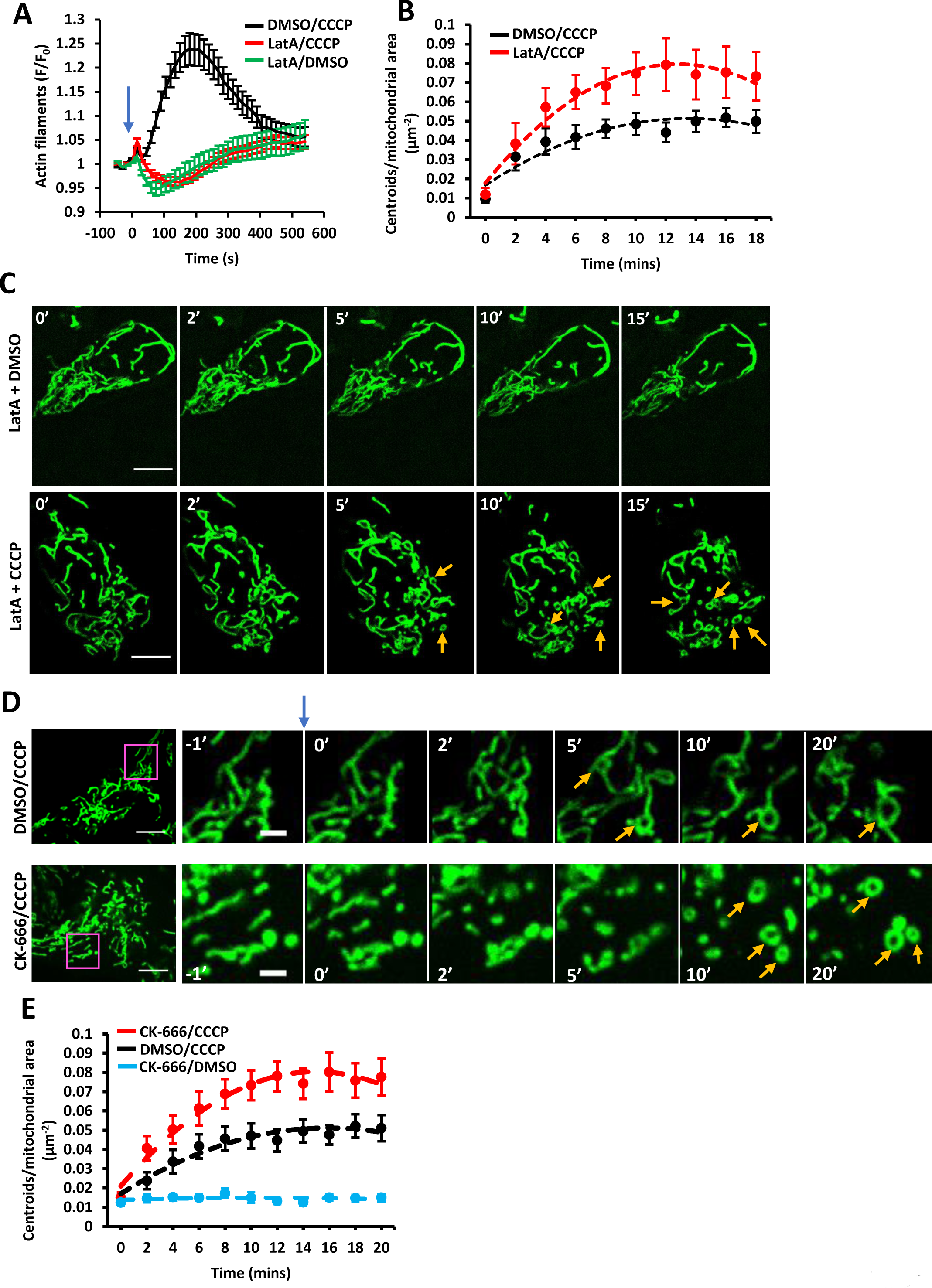
Effect of latrunculin A and Arp2/3 complex inhibition on depolarization-induced mitochondrial shape changes. **(A)** Effect of LatA on CCCP-induced actin polymerization. U2OS cells were treated with combinations of LatA (500 nM) and CCCP (20μM) at time 0 (blue arrow). Confocal images (medial section) were acquired at 15 sec intervals starting four frames before treatment. Data from three experiments. N=23 cells/23 ROIs for LatA/DMSO, 38/38 for LatA/CCCP and 31/31 for DMSO/CCCP. Error bar, ±SEM. **(B)** Effect of LatA on CCCP-induced mitochondrial matrix circularization (‘centroids’). U2OS cells transfected with mito-BFP and GFP-F-tractin. CCCP (20μM) added at time point 0, simultaneous to LatA addition (500 nM). Data from three experiments. N= 17 cells/3702 μm^2^ total mitochondrial area for DMSO/CCCP control cells; 18 cells /3823 μm^2^ for LatA/CCCP. Error bars, ±SEM. **(C)** Time-course montage of LatA-treated cells under control conditions (DMSO treatment, top) or CCCP treatment (20 μM, bottom). U2OS cells, transfected with F-tractin (not shown) and mito-BFP (green), treated with 500nM LatA and CCCP or DMSO simultaneously at 0 min. Confocal images (basal section) acquired at 15 sec intervals starting four frames before treatment. Yellow arrows denote centroids (circular mitochondrial matrix). Time in min. Scale bar: 10μm. **(D)** Time-course montage of CCCP-induced mitochondrial matrix circularization in the absence (top) or presence (bottom) of CK-666 pre-treatment. U2OS cells transfected with GFP-F-tractin (not shown) and mito-BFP (green) were treated with DMSO or CK-666 (100μM) for 30 minutes before stimulation with 20μM CCCP at time point 0 (blue arrow). Confocal images (basal section) were acquired at 15 sec intervals starting four frames before CCCP treatment. Yellow arrows denote centroids (circular mitochondrial matrix). Time in min. Scale bar: 10μm. Inset scale bar: 2.5μm. Corresponds to Movie 9. **(E)** Graph of change in mitochondrial matrix circularization (defined as centroids per total mitochondrial area in the region of interest) from time-courses taken as described in panel A. Data from three experiments. Conditions tested: CK-666 pre-treatment followed by CCCP stimulation (N= 46 cells/8539 μm^2^ total mitochondrial area); DMSO pretreatment, CCCP stimulation (58 cells/11473 total mitochondrial area); and CK-666 pre-treatment/DMSO stimulation (36 cells/6655 total mitochondrial area). Error bars, ±SEM.

We also assessed whether mitochondrial fission was up-regulated during the early stages of CCCP-induced mitochondrial depolarization. For this purpose, we used a live-cell assay to quantify fission rate (Chakrabarti et al., 2018; Ji et al., 2017; Ji et al., 2015), because fixed-cell methods to assess fission based on change in mitochondrial length are confounded by the apparent length change induced by circularization. Our results suggest that CCCP treatment does not increase the number of fission events in the first 30 min, contrary to the fission increase induced by ionomycin (Figure S4A). In addition, we evaluated the effect of Drp1 depletion on CCCP-induced actin clouds and mitochondrial circularization. Drp1 suppression by siRNA has no effect on either of these events, suggesting that acute CCCP induced mitochondrial shape changes are not due to Drp1 dependent mitochondrial dynamics (Fig S4B-D). Combined, these results suggest that the major morphological change to mitochondria in response to depolarization is remodeling of the IMM, resulting in circular mitochondria in which the OMM remains intact.

### Mitochondrial shape changes depend on the IMM protease OMA-1

Since CCCP-induced mitochondrial shape change appears to be due largely to IMM rearrangement, we asked which IMM proteins could be mediating these changes. One candidate is the IMM dynamin family protein Opa1, since Opa1 mediates fusion of IMM (Song et al., 2007), is important for cristae structure (Anand et al., 2014; Patten et al., 2014; Varanita et al., 2015), and also might play a role mitochondrial fission (Anand et al., 2014). Opa1 can be proteolytically processed by two IMM proteases, Oma-1 and Yme-1, with consequences for both mitochondrial fission/fusion balance and for cristae ultrastructure (MacVicar and Langer, 2016). Five distinct bands for Opa1 can be resolved by SDS-PAGE, resulting both from splice variation and differential proteolytic processing (Figure 5A).

**Figure 5:**
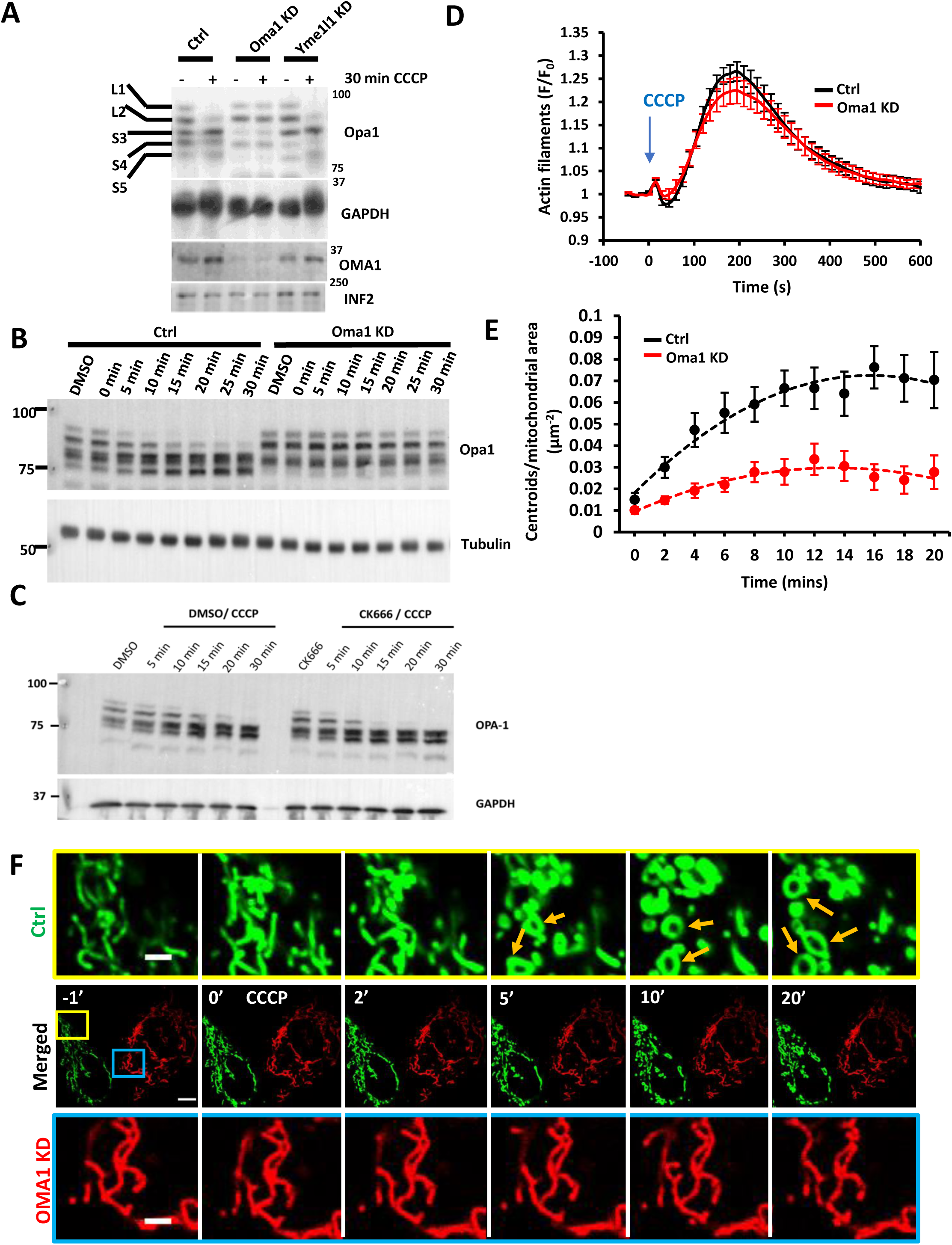
Role of Oma1 in depolarization-induced mitochondrial shape change. **(A)** Western blot of Opa1 and Oma1 in control, Oma1 KD and Yme1 KD U2OS cells before and after treatment with 20μM CCCP for 30 mins. The positions of the five Opa1 forms (L1, L2, S1-3) are indicated. GAPDH, INF2 loading control. **(B)** Western blot of Opa1 in control and Oma1 KD U2OS cells during time course of CCCP treatment (20μM). Tubulin, loading control. **(C)** Western blot of Opa1 in control and CK666 treated U2OS-WT cells during time course of CCCP treatment (20μM). GAPDH, loading control. **(D)** CCCP-induced actin polymerization for control and Oma1 KD U2OS cells, transfected with GFP-F-tractin and mito-BFP, and stimulated with 20μM CCCP (blue arrow). Data from three experiments. N=40 cells/40 ROIs for scrambled control, 28/28 for Oma1 KD. Error bar, ±SEM. **(E)** Change in mitochondrial matrix circularization (‘centroids’) over time for control and Oma1-KD U2OS cells transfected with mito-BFP and GFP-F-tractin. CCCP (20μM) added at time point 0. Data from three experiments. N= 36 cells/4733 μm^2^ total mitochondrial area for control cells; 53 cells /9519 μm^2^ for OMA1 KD cells. Error bars, ±SEM. **(F)** Time-course montage of CCCP-induced mitochondrial shape change (yellow arrows) in control (green) and Oma1-KD U2OS cells (red), imaged in the same field. Control cells (scrambled siRNA) were transfected with Mito-GFP while Oma1-KD cells were transfected with Mito Ds-Red. The two cell populations were trypsinized, mixed, and plated 24 hr before imaging. Confocal images (basal cell section) were acquired at 15 sec intervals starting four frames before CCCP treatment. 20μM CCCP added at time point 0. Time in min. Scale bar: 10μm. Inset scale bar: 2.5μm. Corresponds to Movie 11.

CCCP treatment induces Opa1 proteolytic processing on a similar time scale to both actin polymerization and mitochondrial rearrangement, with increases in short Opa1 bands (S3, S4, S5) within 5 min, and almost complete disappearance of long forms (L1 and L2) by 30 min (Figure 5A, 5B), similar to past results (Ehses et al., 2009; Head et al., 2009; Ishihara et al., 2006). CK666 treatment causes accelerated CCCP-induced Opa1 proteolytic processing (Figure 5C). Conversely, CCCP-induced Opa1 processing is abolished upon siRNA-mediated KD of Oma1 (Figure 5A, 5B), similar to past results (Head et al., 2009; MacVicar and Lane, 2014). In contrast, Yme1 KD has no effect on depolarization-induced Opa1 processing (Figure 5A). Therefore, we reasoned that Oma1 processing of Opa1 might be involved in mediating depolarization-induced mitochondrial shape changes.

We assessed the effect of Oma1-KD or Opa1-KD on CCCP-induced actin polymerization and mitochondrial shape changes. CCCP-induced actin polymerization is unaffected by either Oma1-KD (Figure 5D) or Opa1-KD (Figure S5A, D). Due to the highly fragmented mitochondria resulting from Opa1-KD (Figure S5C), it is difficult to assess the role of Opa1 in CCCP-induced mitochondrial shape change. In contrast, mitochondria in Oma1-KD cells are similar in morphology to those in control cells, allowing examination of shape change upon CCCP treatment. Oma1-KD cells fail to undergo significant CCCP-induced shape changes, as judged by quantifying circularization for 20 min after stimulation (Figure 5E). As a further test of this effect, we mixed control cells (transfected with a GFP-mito marker) and Oma1-KD cells (transfected with a Ds-red mito marker) and imaged the two cell types in the same field upon CCCP treatment. While mitochondria in the control cells undergo CCCP-induced shape changes, mitochondria in the Oma1-KD cells do not (Fig 5F, movie 11).

These results suggest that the rapid changes in mitochondrial morphology induced by CCCP are due to changes in IMM structure through Oma1-mediated proteolysis, rather than through actin-mediated effects on the OMM.

## Discussion

A growing number of studies show that actin can functionally interact with mitochondria (Chakrabarti et al., 2018; Chua et al., 2003; De Vos et al., 2005; DuBoff et al., 2012; Ji et al., 2015; Korobova et al., 2013; Kruppa et al., 2018; Li et al., 2015; Moore et al., 2016; Utsumi et al., 2003). In this paper we show that there are at least two distinct types of mitochondrially-associated actin, differing in cellular stimulus, assembly mechanism, and functional consequences. Mitochondrial depolarization triggers assembly of a dense cloud of actin filaments around the depolarized mitochondria, whereas increased cytosolic calcium triggers actin polymerization throughout the cytosol. Both actin filament populations are transient, with the calcium-induced actin being more rapid both in assembly and disassembly. Depolarization-induced actin is Arp2/3 complex-dependent and INF2-independent, while calcium-induced actin is INF2-dependent and Arp2/3 complex-independent. Inhibition of depolarization-induced actin polymerization causes a greater degree of Oma1-dependent mitochondrial shape change, suggesting that actin polymerization inhibits these changes. Inhibition of calcium-induced actin polymerization causes reduced mitochondrial calcium entry and Drp1 recruitment, with downstream inhibition of mitochondrial fission, which we have shown previously (Chakrabarti et al., 2018; Korobova et al., 2013).

Several studies have shown similar types of actin clouds to those that we observe after depolarization (Kruppa et al., 2018; Li et al., 2015; Moore et al., 2016). In two cases, these clouds were triggered after mitochondrial depolarization (Kruppa et al., 2018; Li et al., 2015), while in one case the clouds assembled without stimulation, were not associated with depolarization, and progressed in a cyclic pattern around the cell (Moore et al., 2016). Both the depolarization-induced and the depolarization-independent clouds were blocked by Arp2/3 complex inhibition (Kruppa et al., 2018; Moore et al., 2016). Therefore, multiple mechanisms may exist to activate Arp2/3 complex around mitochondria. In addition, these studies show formins are involved in Arp2/3 complex-mediated actin assembly around mitochondria (Kruppa et al., 2018; Moore et al., 2016). In our work, we find that INF2 is not the responsible formin in the case of depolarization-induced clouds. Mammals possess 15 distinct formin proteins (Higgs and Peterson, 2005; Pruyne, 2016), and it will be interesting to identify which formin(s) participate(s) in actin cloud assembly.

One question concerns the activation mechanisms for INF2 and Arp2/3 complex in these processes. We recently showed that cytosolic calcium activates INF2 through a mechanism involving HDAC6-mediated deacetylation of actin (A et al., 2019). The mechanism by which mitochondrial depolarization activates Arp2/3 complex is less clear. Arp2/3 complex is directly activated by members of the nucleation-promoting factor (NPF) family, which includes WASP/N-WASP, WAVE proteins, WASH, Dip/WISH, WHAMM, JMY, and cortactin (Campellone and Welch, 2010). The NPF responsible for depolarization-induced Arp2/3 complex activation has yet to be identified. WHAMM is an attractive candidate, being recruited to early autophagosomes to promote Arp2/3 complex activity during starvation-induced non-specific autophagy (Kast et al., 2015). Since long-term CCCP incubation leads to mitophagy and an overall increase in autophagosomes (Li et al., 2018; Matsuda et al., 2010; Park et al., 2018; Tanaka et al., 2010; Vives-Bauza et al., 2010). WHAMM may also regulate actin clouds around mitochondria. JMY is another strong candidate, due to its role in autophagosome formation after cell stress (Coutts and La Thangue, 2015). Finally, previous results show that downregulation of cortactin results in elongated and interconnected mitochondria (Li et al., 2015), so cortactin is also a candidate Arp2/3 complex activator.

Another question is how IMM depolarization can activate the NPF involved in actin cloud assembly. One protein activated by depolarization is the IMM protease Oma1, but we show that neither Oma1 suppression, nor suppression of the Oma1 substrate Opa1, inhibits actin cloud assembly. Another interesting candidate to relay the depolarization signal might be the protein kinase PINK1, which is stabilized on the outer mitochondrial membrane (OMM) of depolarized mitochondria (Narendra et al., 2010). An important role of PINK1 is to initiate PARKIN-mediated mitophagy by phosphorylating both PARKIN and ubiquitin (Okatsu et al., 2013; Pickles et al., 2018; Yamano et al., 2018). PINK1 is expressed in U2OS cells (McLelland et al., 2018) while PARKIN has not been detected (Durcan et al., 2014), so it is possible that stabilized PINK1 phosphorylates other substrates that control Arp2/3 complex activation. Candidates include the recently discovered PINK1 substrate Paris (Lee et al., 2017), or possibly direct phosphorylation of an NPF.

A third question concerns the roles for actin clouds around mitochondria. There is evidence that actin filaments are involved in mitochondrial quality control (Kruppa et al., 2018; Li et al., 2015). A recent paper (Kruppa et al., 2018) showed evidence that “cages” of actin and myosin VI assembled around CCCP-depolarized mitochondria and inhibited mitochondrial fusion after CCCP wash-out. These cages assemble later (approximately 2 hrs) than the actin clouds shown here. Since mitochondrial fusion has been shown to require membrane potential (Hoppins et al., 2011; Legros et al., 2002; Song et al., 2007), and mitochondria remain depolarized for at least 10 min after CCCP treatment in U2OS cells (Figure S1A), it is unlikely that the rapidly-assembled actin clouds act as a mitochondrial fusion barrier.

Another possibility is that actin polymerization mediates changes in mitochondrial morphology, since CCCP treatment results in significant changes in mitochondrial structure on a similar time course as actin cloud assembly. However, we find that these rapid mitochondrial shape changes still occur under conditions that prevent actin cloud assembly. In fact, the shape changes are actually increased in the absence of actin clouds. Possibly, actin clouds serve to confine the depolarized mitochondria in order to reduce mitochondrial dynamics.

Our work provides insights into the mechanisms behind these shape changes. A number of studies suggest that depolarization-induced shape changes are the result of massive mitochondrial fragmentation, which are generally observed after mitochondria have been depolarized for an hour or longer (Duvezin-Caubet et al., 2006; Fu and Lippincott-Schwartz, 2018; Kwon et al., 2017; Li et al., 2018). However, others show that in the first 20 min of depolarization the major change in mitochondria is circularization, which is due to changes in the IMM but not the OMM (De Vos et al., 2005; Liu and Hajnoczky, 2011; Miyazono et al., 2018).

Our live-cell imaging agrees with these observations, showing that the IMM circularizes while the OMM remains intact. IMM circularization depends upon OMA-1 and is independent of Drp1. We also show that inhibition of Arp2/3 complex increases the rate of OMA-1 mediated OPA1 processing after mitochondrial depolarization. It is therefore possible that signaling occurs in both directions in this system, with mitochondrial depolarization triggering cytosolic Arp2/3 complex activation, and the resulting actin clouds inhibiting OMA-1 activity. Subsequent actin cloud disassembly might then allow OMA-1 to mediate IMM rearrangement. While OPA1 is likely to be the relevant OMA-1 substrate in these rearrangements, other OMA-1 substrates have been identified as well, including PINK1 (Sekine et al., 2019); C11orf83 (Desmurs et al., 2015) and even OMA-1 itself (Zhang et al., 2014). Reciprocal communication from and to the mitochondrial matrix, through actin polymerization, could serve as another form of interaction between the mitochondrion and its cellular environment.

## Acknowledgements

We thank past and present members of the Higgs lab, especially Lori Schoenfeld for generating the U2OS INF2 KO cell line. We also thank Zdenek Svindrych and Ann Lavanway for microscopy help and valuable tips. Being ever-present, the importance of Uli Toppanos cannot be overstated. This work supported by NIH R35 GM122545, DK088826, and P20 GM113132.

## Supplementary Figures

**Figure S1.**
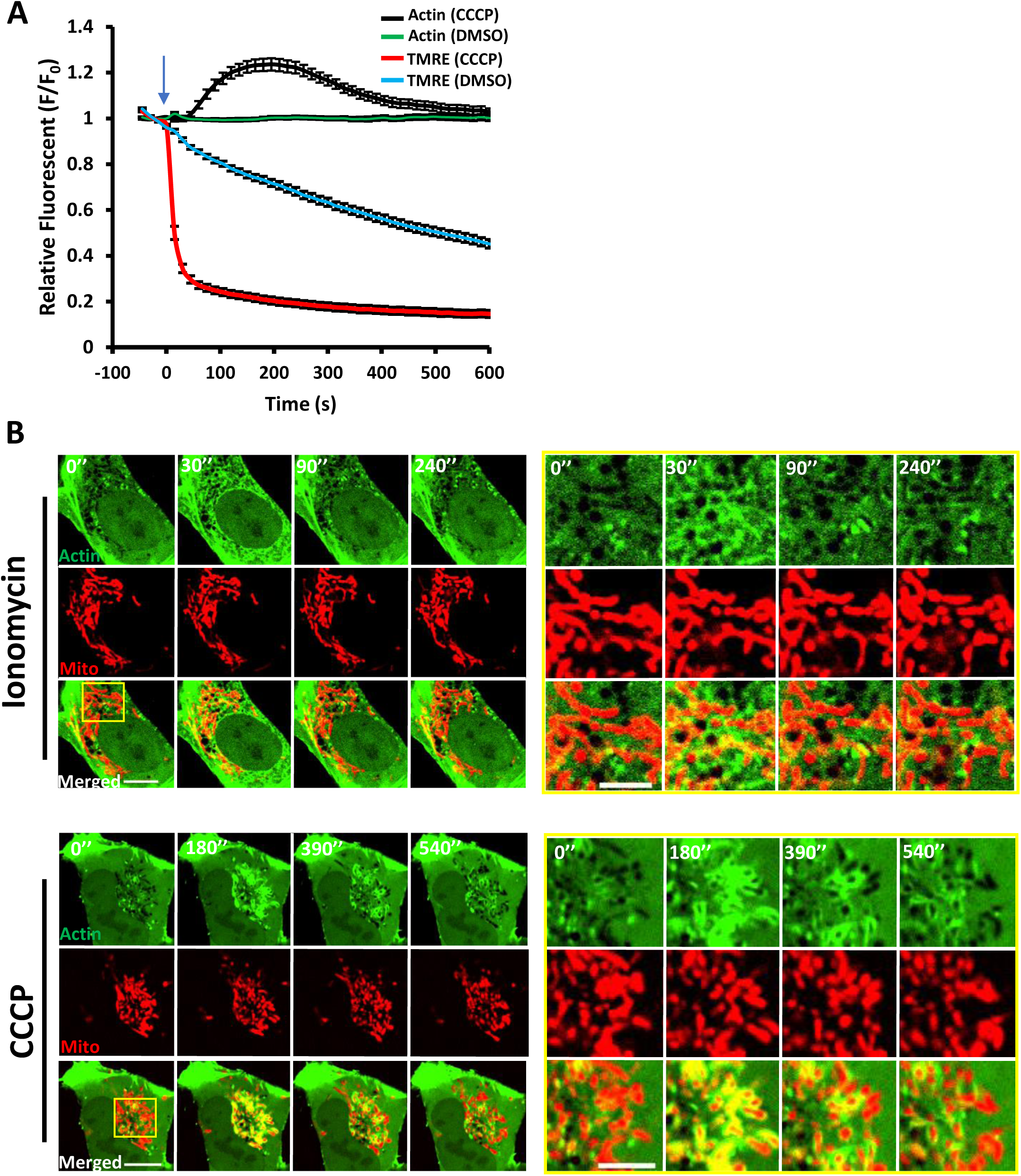
Actin and mitochondrial dynamics upon ionomycin and CCCP treatments. **(A)** CCCP-induced actin polymerization and mitochondrial depolarization in U2OS cells. Cells transfected with GFP-F-tractin and mito-BFP, stained with 20 nM TMRE for 30 min, and stimulated with 20μM CCCP or DMSO at 0 sec (blue arrow). Confocal images (medial section) acquired at 15 sec intervals starting four frames before CCCP treatment. F-tractin and TMRE intensity quantified. Data from three experiments. N=40 cells/40 ROIs for CCCP, 33/33 for DMSO. Error bar, ±SEM. **(B)** Time-lapse image montage of ionomycin-induced (top) and CCCP-induced (bottom) actin polymerization for U2OS cells transfected with GFP-F-tractin (green) and mito-BFP (red). Imaging conducted at a medial cell section. Ionomycin or CCCP added at time point 0. Scale bar: 10μm. Inset scale bar: 5μm. Corresponds to Movies 3 and 4.

**Figure S2.**
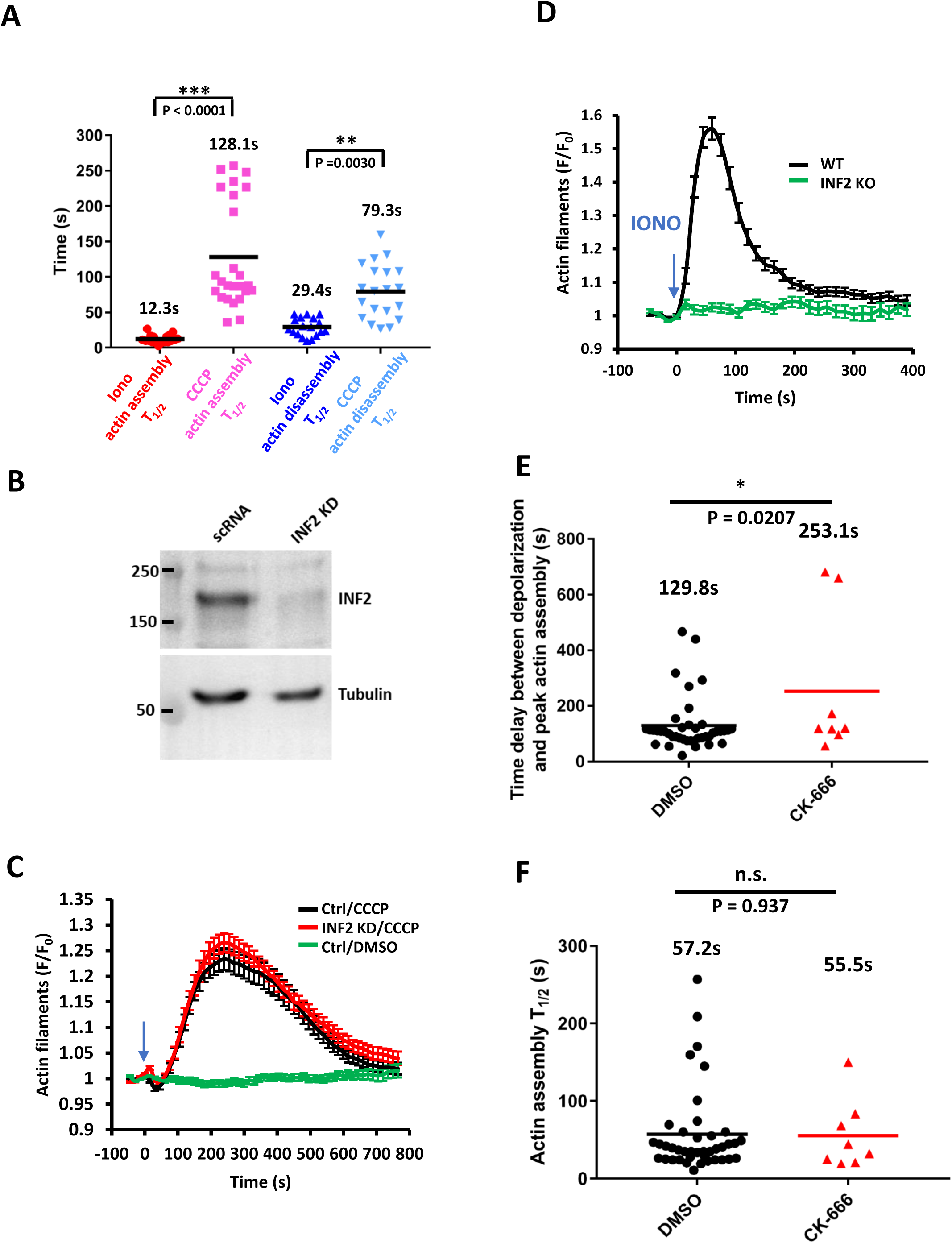
Actin dynamics upon CCCP treatment in control and INF2 KD cells. **(A)** Scatter plots of actin stimulation and recovery half-times for ionomycin and CCCP treatments in U2OS-WT cells imaged at 1.4 sec/frame. Number of individual cells: 18 (ionomycin) and 25 (CCCP/ actin assembly); 20 (CCCP actin disassembly). Data compiled from three independent experiments. P value (unpaired) using Sidak’s multiple comparisons test. Error, Standard deviation. **(B)** Western blot for INF2 of control (scrambled siRNA) and INF2-KD cells. Tubulin, loading control. **(C)** Graph of CCCP-induced actin polymerization for control (scrambled siRNA) and INF2-KD U2OS cells. Cells were transfected with GFP-F-tractin and mito-BFP, then stimulated with DMSO or 20μM CCCP (blue arrow). Confocal images (medial section) acquired at 15 sec intervals starting four frames before CCCP treatment. Data from three experiments. N= 67 cells/67 ROIs for control/CCCP, 53/53 for INF2 KD/CCCP, and 26/26 for control/DMSO. Error bar, ±SEM. **(D)** Graph of ionomycin-induced actin polymerization for U2OS-WT and INF2-KO cells. Cells transfected with GFP-F-tractin and mito-BFP, then stimulated with 4μM ionomycin (blue arrow). Confocal images (medial section) acquired at 15 sec intervals starting four frames before ionomycin treatment. Data from three experiments. N = 30 cells/60 ROIs for WT cells, 27/54 for INF2 KO cells. Error bar, ±SEM. **(E)** Lag between mitochondrial depolarization and actin assembly during spontaneous depolarization events. From the same data set described in Figure 2. Student’s unpaired t-test. **(F)** Scatter plots of actin stimulation during spontaneous depolarization events. U2OS cells transfected with GFP-F-tractin and mito-BFP, stained with 20 nM TMRE for 30 min, then imaged by confocal microscopy for 20 min. From the same data set described in Figure 2. Student’s unpaired t-test.

**Figure S3.**
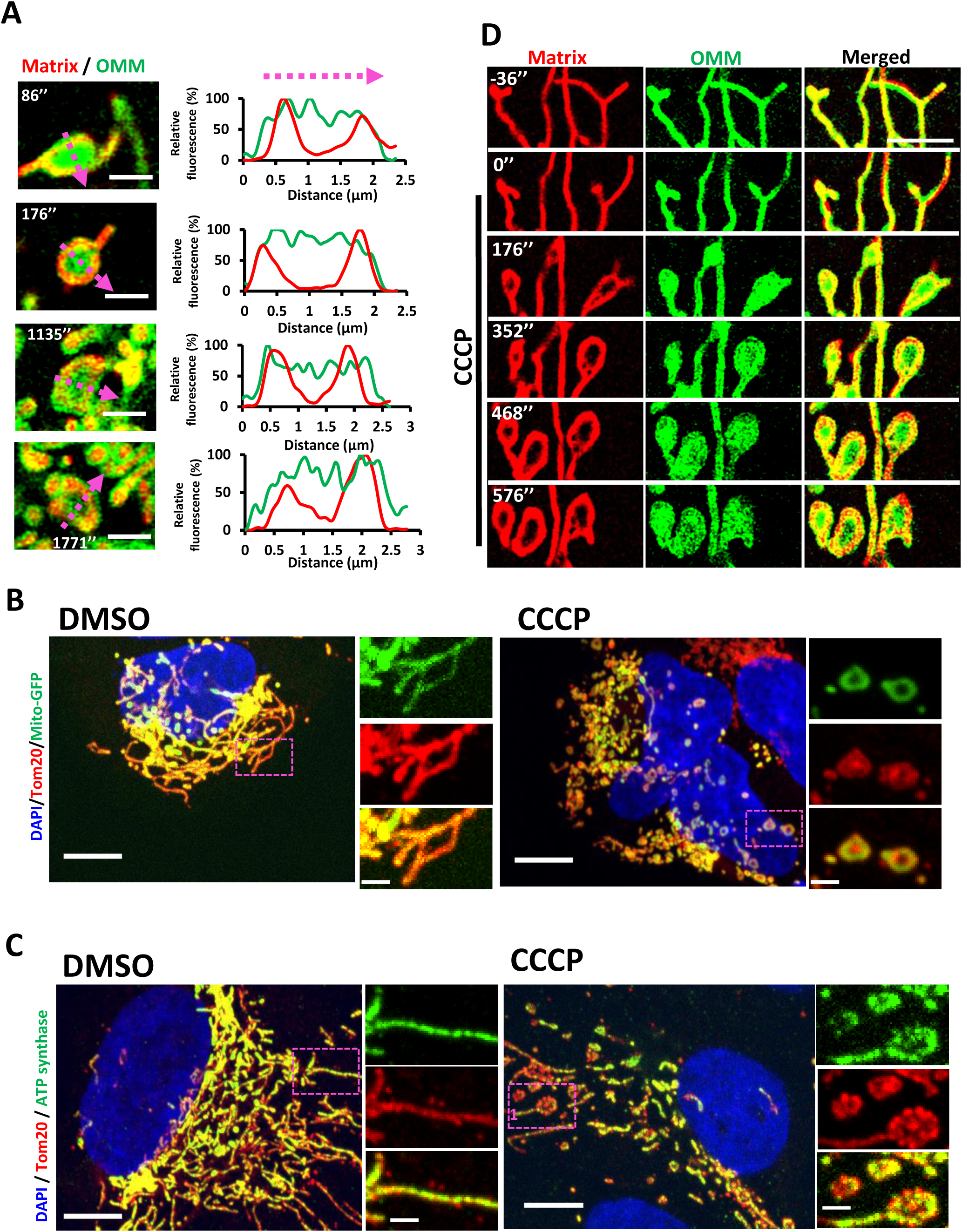
Depolarized mitochondria with circular matrix and intact OMM. **(A)** Four examples of CCCP-induced mitochondrial shape changes. U2OS cells transfected with Mito-DsRed (matrix, red) and Tom20-GFP (OMM, green) and stimulated with 20μM CCCP at time point 0. Airyscan microscopy images (basal cell section) acquired at 4.2 sec intervals starting ten frames before CCCP treatment. Single time points at time of matrix circularization shown (time (sec) after CCCP addition shown in upper left). Scale bar: 2μm. On right, line scans for matrix and OMM taken along the magenta line, showing reduced signal for the matrix marker but not for OMM marker. **(B)** Maximum intensity projections of paraformaldehyde-fixed U2OS cells (expressing the matrix marker mito-GFP, green) after treatment with either DMSO (top) or 20μM CCCP (bottom) for 20 min. Cells were stained for OMM using anti-Tom20 (red) and nucleus using DAPI (blue). Z stacks were taken at step size of 0.4μm. Representative examples of mitochondrial circularization after CCCP treatment are zoomed in. Scale bars: 10μm and 2μm (insets). **(C)** Maximum intensity projections of paraformaldehyde fixed U2OS cells after treatment with either DMSO (top) or 20μM CCCP (bottom) for 20 min. Cells were stained for the IMM with ATP synthase (green), OMM with anti-Tom20 (red) and DAPI (blue) for nucleus. Z stacks were taken at step size of 0.4μm. Representative examples of mitochondrial circularization after CCCP treatment are zoomed in. Scale bar: 10μm and inset 2μm. **(D)** Dynamics of the OMM and mitochondrial matrix upon CCCP treatment in INF2-KO cells. INF2-KO U2OS cell transfected with Mito-R-GECO1 (red) and Tom20-GFP (green) was treated with 20μM CCCP at time point 0. Airyscan images (basal region) acquired at 4 sec intervals starting 10 frames before CCCP treatment. Scale bar: 5μm. Corresponds to Movie 10.

**Figure S4:**
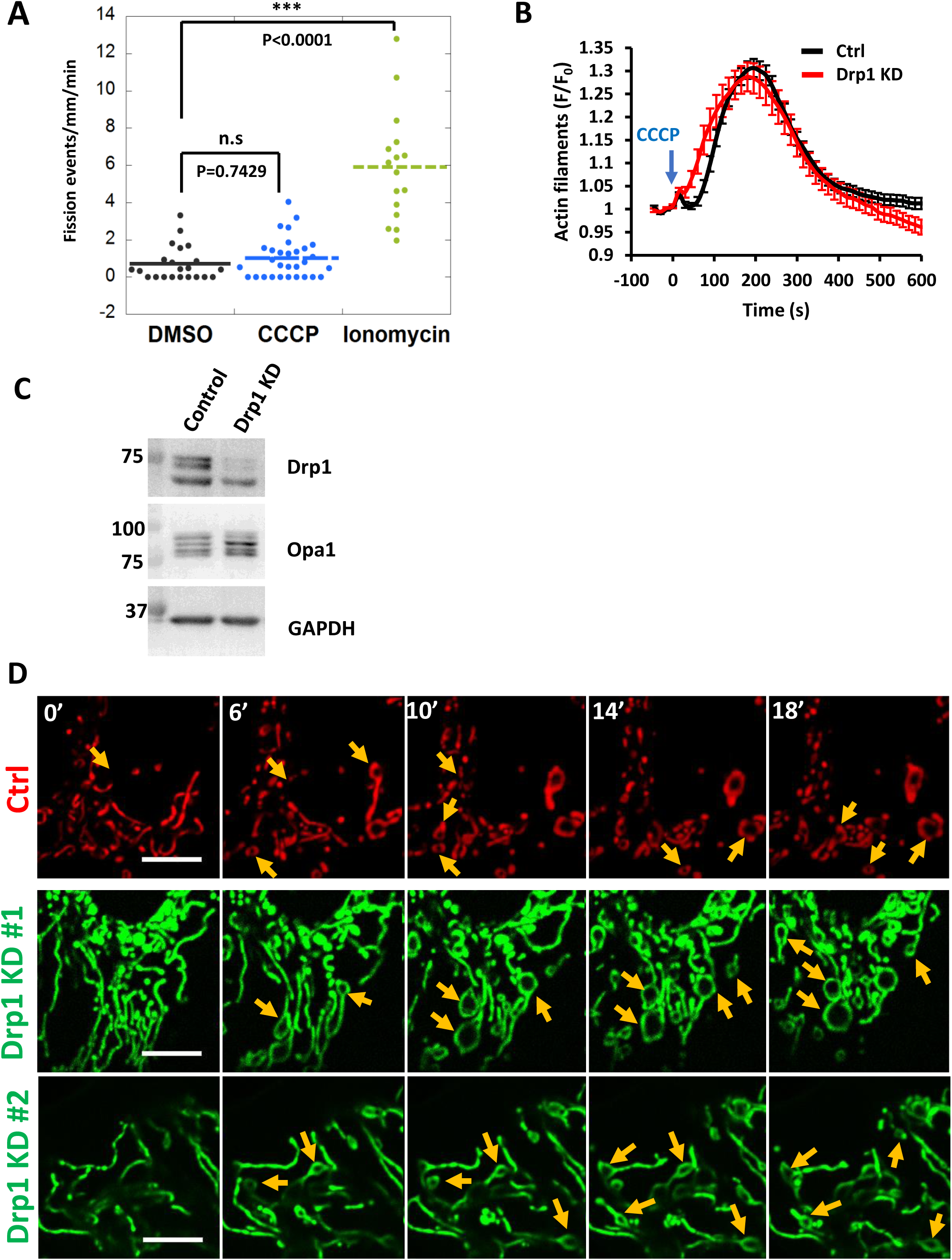
Effect of Drp1 KD on CCCP induced actin clouds and mitochondrial shape change. **(A)** Quantification of mitochondrial fission rate in U2OS cells, using ROIs from live-cell movies of mitochondrial matrix marker (mito-BFP) treated with DMSO, 4 µM ionomycin (10 mins) or 20µM CCCP (30 min). N = 22 cells, 2590.5 µm mitochondrial perimeter (DMSO), 16 cells, 3093.7 µm mitochondrial perimeter (ionomycin), 30 cells, 3999.5 µm mitochondrial perimeter (CCCP). Each point represents one ROI per cell. Compiled from two independent experiments. Dunnett’s multiple comparisons test (unpaired): DMSO vs CCCP, p=0.7429 and DMSO vs Ionomycin, p <0.0001 **(B)** Effect of Drp1 depletion on CCCP-induced actin polymerization. CCCP-induced actin polymerization for control and Drp1-KD U2OS cells, transfected with GFP-F-tracin and mito-BFP, stimulated with 20μM CCCP (blue arrow) and imaged at 15 sec intervals. Data from three experiments. N=40 cells/40 ROIs for scrambled control, 41/41 for Drp1 KD. Error bar, ±SEM. **(C)** Western blot analysis of Drp1 and Opa1 in control (scrambled siRNA) and Drp1 KD U2OS cells. GAPDH, loading control. **(D)** CCCP induced mitochondrial shape changes in Drp1 KD cells. Time-lapse image montage of CCCP-induced mitochondrial shape change in control (top, red) and Drp1 KD cells (middle, bottom, green) transfected with mito-BFP. Imaging conducted at a basal cell section. CCCP added at time point 0. Scale bar: 10μm.

**Figure S5.**
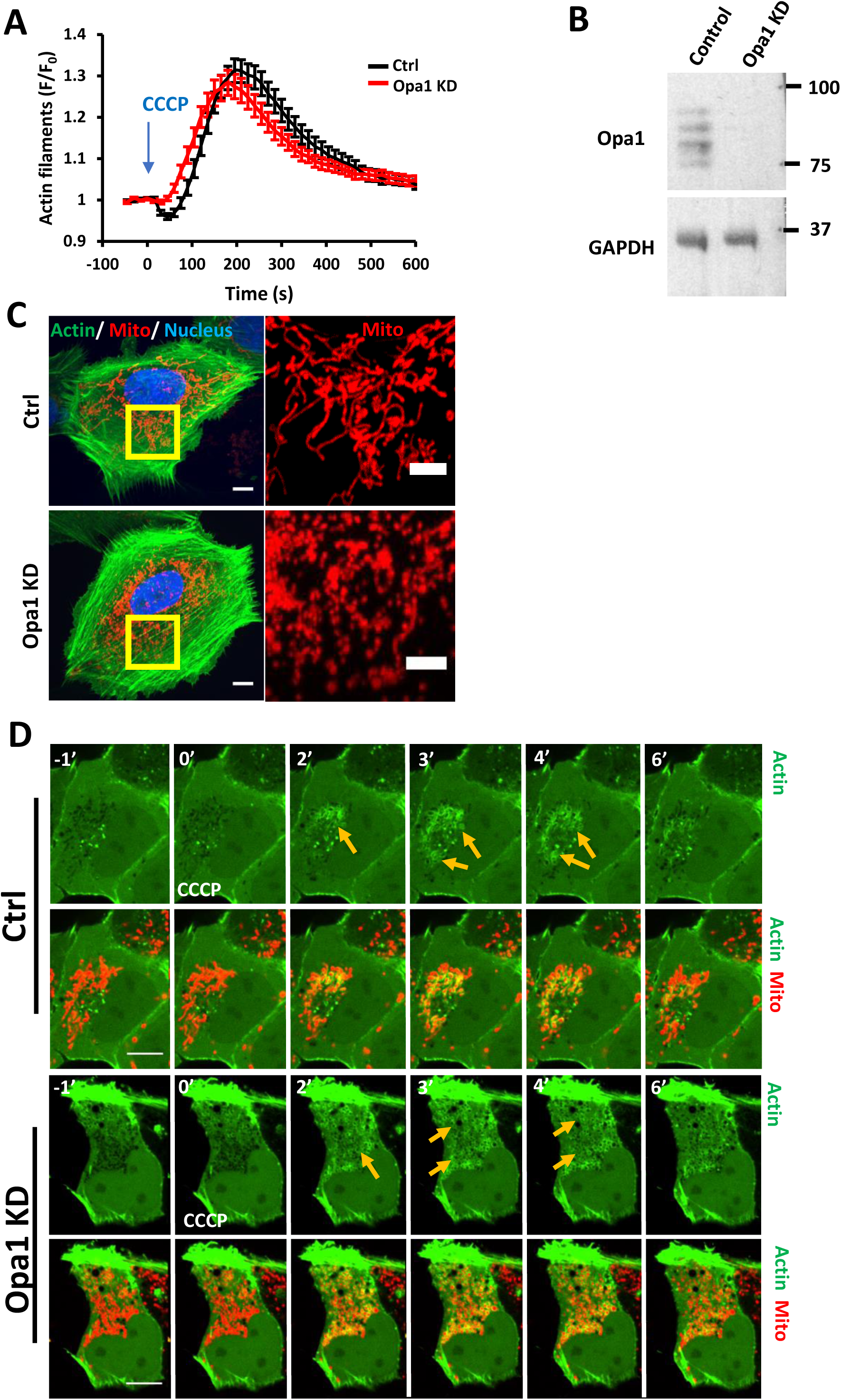
Effect of Opa1 KD on CCCP-induced actin burst and mitochondrial morphology. **(A)** CCCP-induced actin polymerization for control and Opa1-KD U2OS cells, transfected with GFP-F-tracin and mito-BFP, stimulated with 20μM CCCP (blue arrow) and imaged at 15 sec intervals. Data from three experiments. N=23 cells/23 ROIs for scrambled control, 27/27 for Opa1 KD. Error bar, ±SEM. **(B)** Western blot analysis of Opa1 in control (scrambled siRNA) and Opa1 KD U2OS cells. GAPDH, loading control. **(C)** Maximum intensity projection of control U2OS cells (scrambled siRNA) and Opa1 KD U2OS cells transfected GFP-Ftractin, mitoBFP and H2B-mCherry. Z sections were selected based on actin and imaged at 0.4 µm stepsize. Scale bar: 10μm and inset scale bar: 5μm. **(D)** Time-course of CCCP induced actin polymerization for control and Opa1 KD U2OS cells transfected with GFP-F-tractin (green) and mito-BFP (red). Confocal images from from a medial cell section. CCCP added at time point 0. Time in minutes. Scale bar: 10μm. Yellow arrows denote actin assembly around mitochondria.

## Movie legends

**Movie 1: Ionomycin-induced actin polymerization in U2OS cells – basal region.**

U2OS cell expressing Ftractin-GFP (green) and mito-BFP (red), stimulated with 4μM Ionomycin at time 0 (sec). Confocal images acquired every 15sec at basal region. Box denotes the region in inset at right. Corresponds to Figure 1A. Scale bar: 10μm for whole cell and 5μm for inset.

**Movie 2: CCCP-induced actin polymerization in U2OS cells – basal region.**

U2OS cell expressing Ftractin-GFP (green) and mito-BFP (red), stimulated with 20μM CCCP at time 0 (sec). Confocal images acquired every 15sec at basal region. Box denotes the region in inset at right. Corresponds to Figure 1A. Scale bar: 10μm for whole cell and 5μm for inset.

**Movie 3: Ionomycin-induced actin polymerization in U2OS cells – medial region.**

U2OS cell expressing Ftractin-GFP (green) and mito-BFP (red), stimulated with 4μM Ionomycin at time 0 (sec). Confocal images acquired every 15sec at medial region (2-4 μm above basal surface). Box denotes the region in inset at right. Corresponds to Figure S1B. Scale bar: 10μm for whole cell and 5μm for inset.

**Movie 4: CCCP-induced actin polymerization in U2OS cells – medial region.**

U2OS cell expressing Ftractin-GFP (green) and mito-BFP (red), stimulated with 20μM CCCP at time 0 (sec). Confocal images acquired every 15sec at medial region 2-4 μm above basal surface). Box denotes the region in inset at right. Corresponds to Figure S1B. Scale bar: 10μm for whole cell and 5μm for inset.

**Movie 5: Transient depolarization and actin polymerization in U2OS cells.**

U2OS cell, pre-treated with DMSO for 30 min, co-transfected with Ftractin-GFP (green) and mito-BFP (red), and stained with TMRE (cyan). Transient depolarization occurs at time 0 (sec). Confocal images acquired every 1.2sec in the medial region. * denotes the region in inset at right. Corresponds to Figure 2A. Scale bar: 10μm for whole cell and 5μm for insert. Cell is the DMSO-treated control for the CK-666 treatment in Movie 6.

**Movie 6: Transient depolarization and actin polymerization in CK-666 pretreated U2OS cells.**

U2OS cell pretreated with 100μM CK-666 for 30 min, co-transfected with Ftractin-GFP (green) and mito-BFP (red) and stained with TMRE (cyan). Transient depolarization occurs at time 0 (sec). Confocal images acquired every 1.2sec in the medial region. * denotes the region in inset at right. Corresponds to Figure 2A. Scale bar: 10μm for full cell and 5μm for insert.

**Movie 7: Examples of mitochondrial circularization after CCCP treatment**

WT U2OS cells transfected with Mito-DsRed (red) and stimulated with 20μM CCCP at time 0 (sec). Examples show: (A) splitting at a branch point, (B) splitting at the center, (C) splitting at the end, and (D) curling. Confocal images acquired every 4sec at the basal region. Playback: for A, B and D, 4 sec/frame; for C, 16sec/frame. Corresponds to Figure 3A. Scale bar: 2μm.

**Movie 8: Dynamics of the OMM and the mitochondrial matrix during circularization.**

WT U2OS cell co-transfected with Tom20-GFP (green) and Mito-Ds-Red (red), and stimulated with 20μM CCCP at time 0 (sec). Airyscan images acquired every 4sec at the basal region. Corresponds to Figure 3B. Scale bar: 2.5μm.

**Movie 9: Comparison of CCCP-induced circularization in DMSO-pretreated and CK-666-pretreated U2OS cells**

U2OS cells transfected with Ftractin-GFP (not shown) and Mito-BFP (green), and pre-treated with DMSO (left) or 100μM CK-666 (right), then stimulated with 20μM CCCP at 00:00 min:sec. Confocal images acquired every 15sec at the basal region. Corresponds to Figure 4D. Scale bar: 10μm.

**Movie 10: Dynamics of the OMM and the mitochondrial matrix during circularization in INF2-KO cells.**

INF2-KO U2OS cell co-transfected with Tom20-GFP (green) and Mito-R-GECO1 (red) and stimulated with 20μM CCCP at time 0 (sec). Airyscan images acquired every 4sec at the basal region. Corresponds to Figure S3D. Scale bar: 5μm.

**Movie 11: Comparison of CCCP-induced circularization in control and Oma1 KD U2OS cells**

Control cells (scrambled siRNA) were transfected with mito-Ds-Red (green), and Oma1 KD were transfected with mito-BFP (red). At 5 hr after transfection, both types of cell were re-plated on the same coverslip and allowed to adhere for 18 hours prior to imaging. A field containing both red and green cells was located, and confocal images acquired every 15sec. 20μM CCCP was added at time 00:00 min:sec. Corresponds to Figure 5F. Scale bar: 10μm.

## Materials and Methods

### Cell culture

Wild Type (WT) and INF2-KO Human osteosarcoma U2OS cells were grown in DMEM (Corning, 10-013-CV) supplemented with 10% newborn calf serum (Hyclone, SH30118.03) at 37°C with 5% CO_2_. Cell lines were routinely tested negative for mycoplasma contamination using Universal Mycoplasma detection kit (ATCC, 30-1012K) or LookOut Mycoplasma Detection Kit (Sigma-Aldrich, MP0035). The INF2-KO U2OS cell line made by CRISPR-Cas9 is described elsewhere (Chakrabarti et al., 2018).

### DNA transfections

For plasmid transfections, cells were seeded at 4×10^5^ cells per well in a 35 mm dish ∼ 16 hours before transfection. Transfections were performed in OPTI-MEM media (Gibco, 31985062) with 2μl Lipofectamine 2000 (Invitrogen, 11668) per well for 6 hours, followed by trypsinization and re-plating onto glass-bottomed dishes (MatTek Corporation, P35G-1.5-14-C) at ∼3.5×10^5^ cells per well. Cells were imaged ∼16-24 hours after transfection.

The following expression constructs were used: Mito-DsRed and mito-BFP, previously described (Korobova et al., 2014), consisting of amino acids 1-22 of *S.cerevisiae* COX4 N-terminal to the respective fusion protein. GFP-F-tractin was a gift from Clare Waterman and Ana Pasapera (NIH, Bethseda, MD) and described in (Johnson and Schell, 2009). GFP-Mito was purchased from Clontech (pAcGFP1-Mito, #632432) and consists of the mitochondrial targeting sequence derived from the precursor of subunit VIII of human cytochrome c oxidase. Tom20-GFP was made by restriction digest of Tom20 from Tom20-mCherry (a gift from Andrew G York, NIH, Bethseda, MD) with NheI and BamHI; and cloned into eGFP-N1 (Clontech) and is previously described in (Chakrabarti et al., 2018). Mito-R-GECO1 (Addgene, #46021) is previously described in (Wu et al., 2014). H2B-mCherry (Addgene # 20972) is previously described in (Nam and Benezra, 2009).

The following amounts of DNA were transfected per well (individually or combined for cotransfection): 500ng for Mito-BFP, Mito-DsRed, GFP-Mito, Mito-R-GECO1, H2B-mCherry and GFP-F-tractin and 600ng for Tom20-GFP construct.

For siRNA transfections, 10^5^ cells were plated on a 35 mm dish and 2μl RNAimax (Invitrogen, 13778) with 63pg siRNA were used per well. Cells were analyzed 96 hours post siRNA transfection. For live-cell imaging, plasmids containing fluorescent markers were transfected into siRNA-treated cells 18-24 hrs prior to imaging, as described above. All siRNAs were purchased from IDT Inc, including: human INF2 (custom synthesized, HSS.RNAI.N001031714.12.7., 5’-GGAUCAACCUGGAGAUCAUCCGC-3’); human Oma1 (hs.Ri.OMA1.13.1, 5’-GGAUAUUCAGGGUCAAAUGUACAUGAUUUGACCCUG-3’); human Yme1l1 (hs.Ri.YME1L1.13.1, 5’-GGUGGAGGAAGCUAAACAAGAAUUA-3’); human Opa1 (hs.Ri.OPA1.13.1, 5’-CCACAGUGGAUAUCAAGCUUAAACA-3’); and negative control (#51-01-14-04, 5’-CGUUAAUCGCGUAUAAUACGCGUAU-3’).

### Antibodies

Anti-INF2 (rabbit polyclonal against amino acids 941-1249 of human INF2) was described in (Ramabhadran et al., 2011) and used at 3.75 μg/mL. Anti-Opa1 (BD Biosciences, 612606, mouse monoclonal, clone 18/OPA1) was used at 1:2000. Anti-Oma1 (Santa Cruz Biotechnology, sc-515788, mouse monoclonal, clone H-11/OMA1) was used at 1:500. Anti-tubulin (Sigma-Aldrich, T9026, mouse, clone DM1-α) was used at 1:10,000 dilution. Anti-GAPDH (Santa Cruz

Biotechnology, sc-365062, G-9, mouse) was used at 1:1500. Anti-Tom20 (Abcam, ab78547) was used at 1:500 for immunofluorescence. Anti-ATP synthase beta monoclonal antibody (Invitrogen, A-21351, mouse, 3D5AB1) was used at 1:500 for immunofluorescence. Secondary antibodies used for westerns were: goat anti-mouse IgG horseradish peroxidase (HRP) conjugate (Bio-rad, 1705047) at 1:2000 and goat anti-rabbit IgG HRP conjugate (Bio-rad, 1706515) at 1:5000. For immunofluorescence, we used goat anti-rabbit IgG Texas red secondary (Vector Laboratories, TI-1000) was used at 1:500, and horse anti-mouse IgG Fluorescein secondary (Vector Laboratories, FI-2000) was used at 1:500.

### Western blot analysis

Cells from a 35 mm dish were trypsinized, pelleted by centrifugation at 300xg for 5 min, and resuspended in 400μl 1 X DB (50mM Tris-HCl, pH 6.8, 2mM EDTA, 20% glycerol, 0.8% SDS, 0.02% Bromophenol Blue, 1000mM NaCl and 4M Urea). Proteins were separated by SDS-PAGE in a BioRad mini gel system (7 × 8.4 cm) and transferred onto polyvinylidene fluoride membrane (EMD Millipore, IPFL00010). The membrane was blocked with TBS-T (20mM Tris-HCl, pH 7.6, 136mM NaCl and 0.1% Tween-20) containing 3% BSA (VWR Life Science, VWRV0332) for 1 hour, then incubated with primary antibody solution at 4°C overnight. After washing with TBS-T, the membrane was incubated with HRP-conjugated secondary antibody for 1 hour at 23°C. Signals were detected by chemiluminescence. For western blots of OPA1, samples were prepared and separated by SDS-PAGE on a Hoefer SE600 (14 cm X 14 cm) apparatus and transferred using a Hoefer transfer apparatus. The rest of the procedure was similar to as listed above.

### Immunofluorescence

U2OS-WT cells (1×10^5^, either transfected with mito-GFP or untransfected) were plated on Mat-Tek dishes (MatTek Corporation, P35G-1.5-14-C) 16 hours prior to fixation and staining. Cells were treated with DMSO or 20 uM CCCP for 20 min at 37°C/5% CO_2_, washed twice in PBS (23°C) and fixed in either 1% glutaraldehyde (EMS, 16020) prepared in BRB80 buffer (80 mM PIPES pH 6.9, 1mM MgCl2, 1mM EGTA) or 4% prewarmed paraformaldehyde (EMS, 15170) in PBS for 10 min or 20 min respectively. The glutaraldehyde-fixed samples were additionally washed with NaBH4 (1mg/ml in PBS; 3 × 10 min each) prior to permeabilization. The cells were then permeabilized in 0.1% Triton X-100 in PBS for 10 min and blocked in PBS + 10% calf serum for 30 min. These cells were then stained with Tom 20 antibody and/or ATP-synthase (mitochondria), appropriate secondary antibodies and DAPI (nucleus) in PBS + 1% calf serum and imaged by Dragonfly 302 spinning disk confocal (Andor Technology) CFI Plan Apochromat Lambda 100X/1.45 NA oil in PBS on the Mat-Tek dish. Z-stacks were taken from the basal region to the apical top at 0.4μm step size. Maximum intensity projections were generated from z-stack images and background subtracted in ImageJ Fiji (rolling ball 20.0).

### Live imaging by confocal microscopy and Airyscan microscopy

All live cell imaging was conducted in DMEM (Gibco, 21063-029) with 25mM D-glucose, 4mM L-glutamine and 25mM Hepes, supplemented with 10% newborn calf serum, hence referred to as “live cell imaging media”. Cells (∼3.5×10^5^) were plated onto MatTek dishes 16hrs prior to imaging. Medium was preequilibrated at 37°C and 5% CO_2_ before use.

For confocal microscopy, dishes were imaged using the Dragonfly 302 spinning disk confocal (Andor Technology) on a Nikon Ti-E base and equipped with an iXon Ultra 888 EMCCD camera, a Zyla 4.2 Mpixel sCMOS camera, and a Tokai Hit stage-top incubator set at 37°C. A solid-state 405 smart diode 100 mW laser, solid state 560 OPSL smart laser 50 mW laser, and solid state 637 OPSL smart laser 140 mW laser were used. Objectives used were the CFI Plan Apochromat Lambda 100X/1.45 NA oil (Nikon, MRD01905) for all drug treatment live-cell assays; and CFI Plan Apochromat 60X/1.4 NA oil (Nikon, MRD01605) to observe transient depolarization events during live-cell imaging. Images were acquired using Fusion software (Andor Technology, version 2.0.0.15). For actin burst and TMRE quantifications, cells were imaged at a single confocal slice at the medial region, approximate 2μm above the basal surface, to avoid stress fibers. To observe mitochondrial morphological changes, cells were imaged at a single confocal slice at the basal surface.

For ionomycin treatments, cells were treated with 4μM ionomycin (Sigma-Aldrich I0634, from 1mM stock in DMSO) at the start of the 5^th^ image frame (∼1 min, time interval set at 15sec) during imaging and continued for another 5-10 mins. INF2 KO cells were used as a negative control (Chakrabarti R et al., J. Cell Biol. 2018). For Carbonyl cyanide 3-chlorophenylhydrazone (CCCP) (Sigma-Aldrich, C2759) treatments, cells were treated with 20μM CCCP (from a 100mM stock in DMSO) at the start of the 5^th^ frame (∼1 min, with time interval set at 14sec or 15sec) during imaging and continued for another 15-20mins. Equal volume DMSO (Invitrogen, D12345) was used as the negative control. For Tetramethylrhodamine ethyl ester perchlorate (TMRE) (Sigma-Aldrich, 87917) staining before CCCP treatment, cells were loaded with 20 nM TMRE (from a 30mM stock in DMSO) for 30 minutes in live cell imaging media. Cells were subsequently washed twice with live-cell media and fresh live-cell media was added prior to imaging. During imaging, cells were treated with 20μM CCCP at the start of the 5^th^ frame (∼1 min, with time interval set at 15sec) and continued for another 15-20mins. As a negative control, equal volume DMSO was added during imaging (in place of CCCP) for cells loaded with 20nM TMRE.

To observe transient depolarization in U2OS cells in the absence of CCCP, cells were loaded with 20nM TMRE (with or without CK-666) for 30 minutes in live cell imaging media at 37°C and 5% CO_2_. CK-666 and TMRE treated cells were rinsed twice with fresh live cell medium and 50μM CK-666 containing live cell medium was added prior to imaging. Single field confocal imaging in the medial region was conducted at 1.2sec time interval and continued for 1000 frames (20mins). To visualize more cells in the field, the 60X/1.4NA objective was used. Equal volume DMSO was used as a negative control in place of CK-666 while retaining TMRE.

For Latrunculin A (LatA) (Millipore Sigma, 428021) coupled with CCCP treatment, cells were treated with live-cell media containing 500nM LatA (from a 1mM stock in DMSO) and 20μM CCCP simultaneously at the start of the 5^th^ frame (1 min, time interval set at 15sec). Imaging was continued for 10-20mins. Cells treated with DMSO (replacing LatA) and 20μM CCCP simultaneously were the positive control; while cells treated with 500nM LatA and DMSO (replacing CCCP) simultaneously were used as the negative control.

For CK-666 (Sigma-Aldrich, SML006) pretreatment before CCCP addition, cells were pretreated with 1mL of live-cell media containing 100μM CK-666 (from a 20mM stock in DMSO) for 30 min before the start of imaging. During imaging, cells were treated with 1mL live cell media containing 40μM CCCP at the start of the 5^th^ frame (1 min, time interval set at 15sec). Imaging was continued for 15-20mins with cells in media containing a final concentration of 20μM CCCP and 50μM CK-666. Control cells were pretreated with equal volume DMSO (replacing CK-666) and stimulated with 20µM CCCP during imaging; and as a negative control, cells were pretreated with 100μM CK-666 and equal volume DMSO (replacing CCCP) was added during imaging.

For CK-666 pretreatment before ionomycin addition, cells were pretreated with 1mL of live-cell media containing 100μM CK-666 for 30 mins. After which, cells were directly taken for imaging. During imaging, cells were treated with 1mL of live cell media containing 8μM ionomycin at the start of the 5^th^ frame (1 min, time interval set at 15sec). Imaging continued for 5-10mins with cells in media containing a final concentration of 4μM ionomycin and 50μM CK-666. Control cells were pretreated with equal volume DMSO (replacing CK-666) and stimulated with 4µM ionomycin during imaging; and as a negative control, cells were pretreated with DMSO (replacing CK-666) and additional DMSO (replacing ionomycin) was added during imaging.

For Airyscan imaging, dishes were imaged on the LSM 880 equipped with a 100X/1.4 NA Apochromat oil objective using the Airyscan detectors (Carl Zeiss Microscopy). The Airyscan uses a 32-channel array of GaAsP detector configured as 0.2 airy units per channel. Cells were imaged with the 488nm laser and Band Pass (BP) 420-480/BP 495-620 filter for GFP, 561nm laser and BP 495-550/Long Pass 570 filter for RFP. For live-cell microscopy, WT U2OS cells were co-transfected with 500ng of Mito-DsRed and 600ng Tom20-GFP while INF2 KO cells were co-transfected with 500ng Mito-R-GECO1 and 600ng Tom20-GFP. All imaging conducted at 37°C and 5% CO_2_, with a single basal slice acquired at a frame interval of 4.1sec. Images were subsequently processed using Zen2 software. Raw data were processed using Airyscan processing with Zen Black software (Carl Zeiss, version 2.3).

For mixing experiment, control cells were transfected with 500ng Mito-Ds-Red and Oma1 KD cells were transfected with 500ng Mito-BFP 72 hours post knock down. After four hours transfection, control cells were mixed with Oma1 KD cells in a 1:2 (control: Oma1 KD) volume ratio. Mixed cells were re-plated onto MatTek dishes at ∼3.5×10^5^ cells per dish and allowed to adhere for ∼18 hours. During live-cell imaging with CCCP, fields were selected with both Oma1 KD and control cells visible.

### Quantification from live-cell imaging experiments

Unless otherwise stated, all image analysis was performed on ImageJ Fiji (version 1.51n, National Institutes of Health). Cells that shrunk during imaging or exhibited signs of phototoxicity like blebbing or vacuolization were excluded from analysis.

#### Actin burst measurements

Mean actin fluorescence was calculated by selecting two region of interests (ROIs) (for ionomycin treatments) per cell or one ROI per cell (for CCCP treatments). The ROI selected for CCCP encompass the entire area at the height of actin assembly after CCCP treatment. Fluorescence values for each time point (F) were normalized with the mean initial fluorescence before drug treatment (first four frames – F_0_) and plotted against time as F/F_0_. For DMSO control or cells that did not exhibit actin burst, ROI was selected as the bulk region of the cytoplasm containing mitochondria using the Mito-BFP channel.

#### Centroid measurements

Every eight frame was analyzed (2 min intervals). Imaging fields were coded and scrambled by one investigator (TSF) and given to the other investigator (RC) for blind analysis. Centroid were counted manually for every time point. To normalize the data, the number of centroids was divided by the total mitochondrial area in the field (μm^2^). The results were then decoded by the first investigator.

#### Mitochondrial division rate measurements

Mitochondrial division rate was described in detail previously (Ji et al., 2015). Suitable ROIs were selected based on whether individual mitochondria were resolvable and did not leave the focal plane. One ROI was selected per cell. Files of the ROIs were coded and scrambled by one investigator (RC) and analyzed for division by a second investigator (WKJ) in a blinded manner. The second investigator scanned the ROIs frame by frame manually for division events and determined total mitochondrial length in the ROI using the ImageJ macro, Mitochondrial Morphology. The results were then returned to the first investigator for decoding.

#### Depolarization measurements

Mean TMRE fluorescence was calculated from the entire mitochondrial area determined in the Mito-BFP channel, for which the fluorescent intensity did not change appreciably during imaging. TMRE fluorescence values for each time point (F) were normalized with the mean initial fluorescence before drug treatment (first four frames – F_0_) and plotted against time as F/F_0_.

Transient depolarization events in untreated cells are defined as an abrupt loss in mitochondrial TMRE fluorescence signal that persist for ≥ 30 seconds without an appreciable decrease in fluorescence signal of the Mito-BFP marker. Individual transient depolarization events were manually identified by scrolling through images frame by frame. Clear increase in actin fluorescence signal after transient depolarization were considered as “Δψm followed by actin assembly”; while depolarization events that demonstrated no appreciable actin fluorescence increase were considered as “Δ_ψm_ followed by no actin assembly”. To determine the frequency of depolarization, the total number of depolarization events was normalized to the total cell count and the total duration of the imaging (min). For depolarization duration, suitable ROIs were selected for mitochondria which underwent depolarization. TMRE fluorescence values for each time point were normalized to the mean initial fluorescence (ten frames before transient depolarization) and plotted against time. Depolarization events that occurred 16 minutes into imaging and did not recover at the end of 20 minutes imaging were separately noted.

#### T_1/2_ analysis of actin assembly and disassembly

Mean actin fluorescence was calculated by selecting one region of interest (ROI) (for ionomycin and CCCP treatments). The ROI selected for CCCP encompass the entire cell area at a medial cell section, whereas for ionomycin treatment one ROI at peri-nuclear region (free from stress fibers for all time frame) per cell was selected. Fluorescence values for each time point (F) were normalized with the mean initial fluorescence before drug treatment (first thirty frames – F_0_) and plotted against time as F/F_0_. Half-max value was calculated after establishing the peak value and the time determined from the ascending slope (actin assembly) for each cell. For actin disassembly, the time (s) was determined for the half max value from the descending slope and deducted from the peak time. For ionomycin analysis all the cells were used for assembly, peak and disassembly calculations. For CCCP. 5 cells were removed for disassembly calculations because of stress fiber interference. Number of individual cells: 18 (ionomycin) and 25 (CCCP/ actin assembly & actin peak); 20 (CCCP actin disassembly). Data compiled from three independent experiments.

### Statistical analysis and graph plotting softwares

All statistical analyses and p-value determination were conducted using GraphPad Prism QuickCalcs or GraphPad Prism 7 (version 7.05, GraphPad Software). To determine p-values, an unpaired Student’s t test was performed between two groups of data, comparing full datasets stated in the figure legends. For p-values in multiple comparisons (unpaired), Sidak’s multiple comparisons test and Dunnett’s multiple comparison test were performed in GraphPad Prism 7. All graphs, along with their standard error of mean (SEM) were plotted using Microsoft Excel for Office 365 (version 16.0.11231.20164, Microsoft Corporation), with the exception of scatter plots. Fig 2E, S2A, S2E, S2F were plotted with GraphPad Prism 7 and Fig S4A was plotted with KaleidaGraph (version 4.01, Synergy Software).

